# Whole-Proteome Screening and Multi-Modal Profiling of Antigen-Specific CD4+ T Cells at Single-Cell Resolution

**DOI:** 10.1101/2025.04.21.649828

**Authors:** Rongyu Zhang, Jingqi Qi, Michaela McKasson, Jongchan Choi, Vanessa Gutierrez, Conor Brennan, Sunga Hong, William Chour, Rachel H. Ng, Jingyi Xie, Dan Yuan, Andrew Webster, Simranjeet K. Sidhu, Abby Anderson, Daniel Chen, Rick Edmark, Kim M. Murray, Sarah Li, Connor McDonald, Lee Rowen, Shuo Wang, Yusuf Rasheed, Yapeng Su, Jamie R. Wagner, Jia Chen, Karla Narwaly, Jie Fu, Alexandria Duven, Stephen J. Forman, Mihae Song, Saul Priceman, Christine E. Brown, Antoni Ribas, Deborah Wong, Kelly G. Paulson, Charles W. Drescher, Cristina Puig-Saus, Jason D. Goldman, Cornelia L. Trimble, James R. Heath

## Abstract

Systematic whole-proteome screening and comprehensive profiling of antigen-specific CD4+ T cells are crucial for advancing our understanding of CD4+ T cell immunity, yet such efforts remain technically challenging. Here, we present a high-throughput platform that employs large-scale class II single-chain trimer libraries to detect antigen-specific CD4+ T cells, while simultaneously profiling their antigen specificity, TCRα/β sequences, MHC restriction, whole transcriptomes, and patient/timepoint origins at single-cell resolution. We benchmarked SCTs against conventional pMHCs and validated the SCT library-based approach in direct *ex vivo* identification of antigen-specific CD4+ T cells in healthy donors. We then applied the platform to screen the entire SARS-CoV-2 receptor-binding domain in a longitudinal patient cohort, identifying 2,188 antigen-specific CD4+ T cells and revealing key features that define antigen immunogenicity. Extending to cancer, we performed whole-proteome screening of HPV-16 E6/E7 for TCR repertoire profiling in a precancerous cohort, uncovering functional heterogeneity of HPV-specific TCRs. By integrating high-throughput antigen screening with high-dimensional, multi-modal cellular characterization, our approach offers an unprecedented window into CD4+ T cell immunity across diverse disease contexts and empowers the development of new therapies.

## Introduction

CD4+ helper T cells play a critical role in orchestrating and regulating immune responses. Upon exposure to foreign or tumor-derived peptides presented by class II major histocompatibility complex (pMHC), they enhance antigen-presenting cell function, support CD8+ T cell differentiation, and promote B cell activation and antibody maturation^1^. In cancer, CD4+ T cells can help sustain anti-tumor immunity and even exhibit cytotoxic functions^2,3^. A detailed understanding of the dynamic roles of antigen-specific CD4+ T cells is thus crucial for mapping complex immune responses and guiding the development of therapies.

While large-scale screening methods for antigen-specific CD8+ T cells are well-established^4,5,6,7,8,9,10^, analogous platforms for CD4+ T cells are more challenging due to the complexity of class II MHC antigen presentation. High-throughput peptide-pool stimulation followed by activation marker-based sorting captures TCR sequence, but disrupts the native phenotypic state of the cells, which is crucial for fundamental understanding of CD4+ T cell responses^11,12,13,14^. Alternatively, direct *ex vivo* capture of CD4+ T cells using conventional class II pMHC multimers preserves phenotype but is constrained by a combination of the small size of feasible pMHC libraries (n≤15), and the reliance on imperfect computational predictions^15,16,17,18,19,20,21,22^.

Comprehensive profiling of CD4⁺ T cell immunity requires high-throughput approaches capable of capturing and analyzing large, high-dimensional datasets across patient cohorts. Current methods often only capture isolated parameters—such as TCR clonotype, antigen specificity, or MHC restriction—limiting their ability to provide a holistic view of CD4+ T cell responses.

Here we present a high-throughput platform leveraging class II single-chain trimer (SCT) libraries that enables proteome-wide screening for antigen-specific CD4+ T cells, while simultaneously profiling multiple key parameters—including antigen specificity, MHC restriction, whole transcriptome, TCR clonotype, and patient/timepoint origin—at single cell resolution in a single experiment. We validated the performance of SCTs against conventional pMHCs and demonstrated the SCT library-based approach for direct *ex vivo* detection of common pathogen-specific CD4+ T cells in healthy donors. We then applied the platform to screen the entire SARS-CoV-2 spike receptor-binding domain (RBD) in a longitudinal cohort of 22 participants, identifying 2,188 antigen-specific CD4+ T cells. This high-dimensional dataset, capturing six orthogonal modalities at single-cell resolution, revealed class II-restricted immunogenic antigens and documented dynamic phenotypic shifts over the course of infection. To further demonstrate its broad applicability, we extended the platform to cancer and performed whole-proteome screening of HPV-16 oncogenic proteins E6 and E7 in precancerous patients for CD4 TCR repertoire profiling, uncovering functionally heterogeneous TCRs. By integrating high-throughput whole-proteome screening with multi-modal single-cell profiling, our platform offers a comprehensive and scalable framework for studying CD4+ T cell immunity across disease contexts, accelerating discovery of key immunological mechanisms and therapeutic targets.

## Results

### SCT technology development and validation

We previously developed high-throughput SCT platforms for profiling class I-restricted CD8+ T cells responses to viral and tumor neoantigens^7,8,9,10^. Building on this foundation, we designed a modular platform for scalable generation of human class II SCT libraries, extending prior murine constructs^7,23^. The SCT architecture links HLA α, antigen, and HLA β extracellular domains via flexible glycine-rich linkers (L1 and L2), with the antigen encoded as an integral component of the construct, enabling expression as a single construct and supporting high production efficiency (**Fig. 1a**). A partial invariant chain (pIi) enhances SCT stability, while the linker configuration maintains structural integrity^23,24^. By genetically encoding the peptide, our approach eliminates the need for peptide synthesis, significantly improving throughput. Antigen sequences are assembled from two overlapping primers, automatically designed using a custom in-house algorithm (**Fig. 1b, Methods**), achieving >99% success rate as confirmed by PCR. Conserved regions for pIi and linker sequences allow rapid antigen exchange and allele substitution, enabling scalable library construction across multiple HLAs and antigen sets. High-throughput SCT expression is achieved by transient transfection of human-derived Expi293F cells in multi-well format, followed by stream-lined biotinylation and purification workflows optimized for large-scale production.

**Figure 1.**
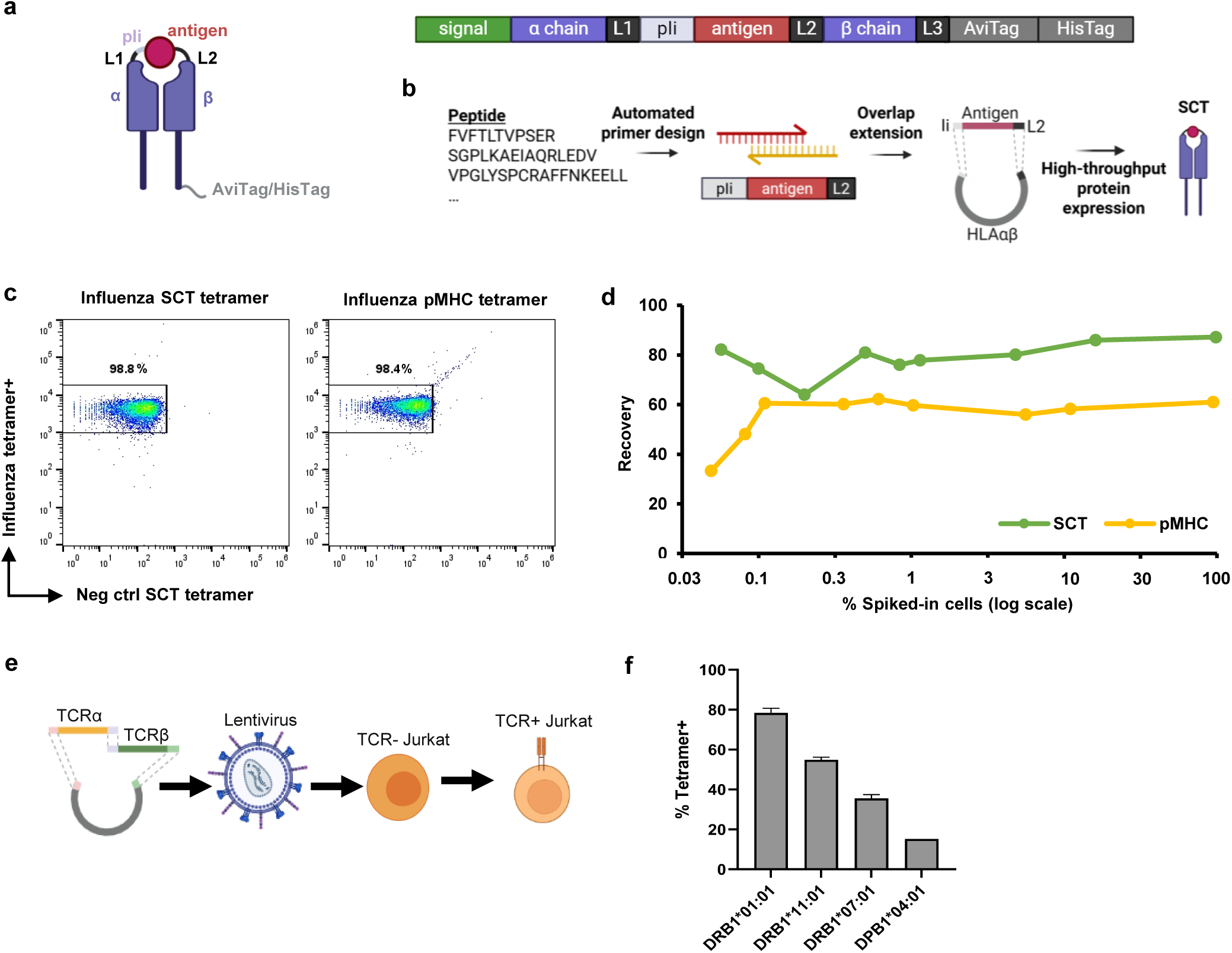
SCT technology development and validation. **a.** Plasmid design encoding the class II SCT protein. The MHC α and β chains are connected to the antigen through flexible linkers (L1, L2( and a partial invariant chain (pli), followed by purification tags (AviTag/His Tag). A secretion signal is placed upstream of the α chain to facilitate protein export. **b.** Experimental workflow for SCT plasmid construction and protein expression. High-throughput production of SCT libraries is enabled by automated primer design and optimized pipeline for plasmid assembly and protein expression/purification. **c.** Comparison of binding efficiency between a class II SCT and a conventional pMHC multimer. Both formats present the DRB1*01:01-restricted influenza peptide (HA_306-318_ PKYVKQNTLKLAT) and were evaluated against influenza-specific CD4+ T cells. An HIV SCT (Gag_41-56_ SALSEGATPQDLNTML) served as a negative control to assess non-specific binding. **d.** Sensitivity comparison netween class II SCTs and conventional pMHC tetramers. **e.** Lentiviral transduction of TCRs into a TCR knock-out Jurkat cell line. **f.** Validation of SCT templates for various common class II HLA alleles through binding to previously reported TCRs.

To benchmark performance, we first validated sensitivity of class II SCTs against conventional pMHCs. A DRB1*01:01-restricted SCT presenting the influenza HA_306-318_ epitope effectively stained a matched influenza-specific CD4+ T cell line, demonstrating binding performance comparable to standard pMHC tetramers with no evidence of non-specific binding from an HIV-specific SCT negative control (**Fig. 1c**). Sensitivity was evaluated by spiking known frequencies of influenza-specific T cells into healthy donor PBMCs. SCT tetramers could detect target cells as low as 0.05% and the recovery rate compares favorably to conventional tetramers (**Fig. 1d**).

We then expanded validation to additional prevalent class II HLA alleles^25,26^, selecting literature-validated TCR-antigen pairs for one HLA-DP and three HLA-DR alleles^27,28,29,30,31^ (**Table S1**). TCRs were cloned into TCR^KO^ Jurkat cells via lentiviral transduction (**Fig. 1e**) and evaluated for binding against their cognate SCTs. Each SCT specifically recognized its matched TCR, with no evidence of cross-reactivity to unrelated SCTs (**Fig. 1f, Extended Data Fig. 1a**). Notably, SCTs displayed post-translational modifications consistent with natural antigen processing, in contrast to synthetic peptide-based approaches (**Extended Data Fig. 1b**). These findings underscore class II SCTs as a robust platform for generating pMHC-like reagents capable of highly sensitive and specific CD4+ T cell detection.

### Benchmarking SCT libraries for antigen-specific CD4+ T cell discovery in healthy donors

Following validation of individual SCTs, we evaluated the performance of SCT libraries in direct *ex vivo* screening for antigen-specific CD4+ T cells to common pathogens in healthy donor peripheral blood mononuclear cells (PBMCs). The modular plasmid design enabled rapid assembly of a 23-elment SCT library presenting DRB1*01:01-restricted canonical antigens from cytomegalovirus (CMV), Epstein-Barr virus (EBV), influenza, and tetanus (collectively, the CEFT library) (**Fig. 2a, Table S2**). PBMCs from five HLA-matched healthy donors were enriched for CD4+ T cells and stained with PE- and APC-labeled SCT tetramers (**Fig. 2b**). Double-positive cells were sorted for single-cell sequencing of paired TCRα/β chains, while cells binding to negative control tetramers were excluded (**Fig. 2c, Extended Data Fig. 1c**).

**Figure 2.**
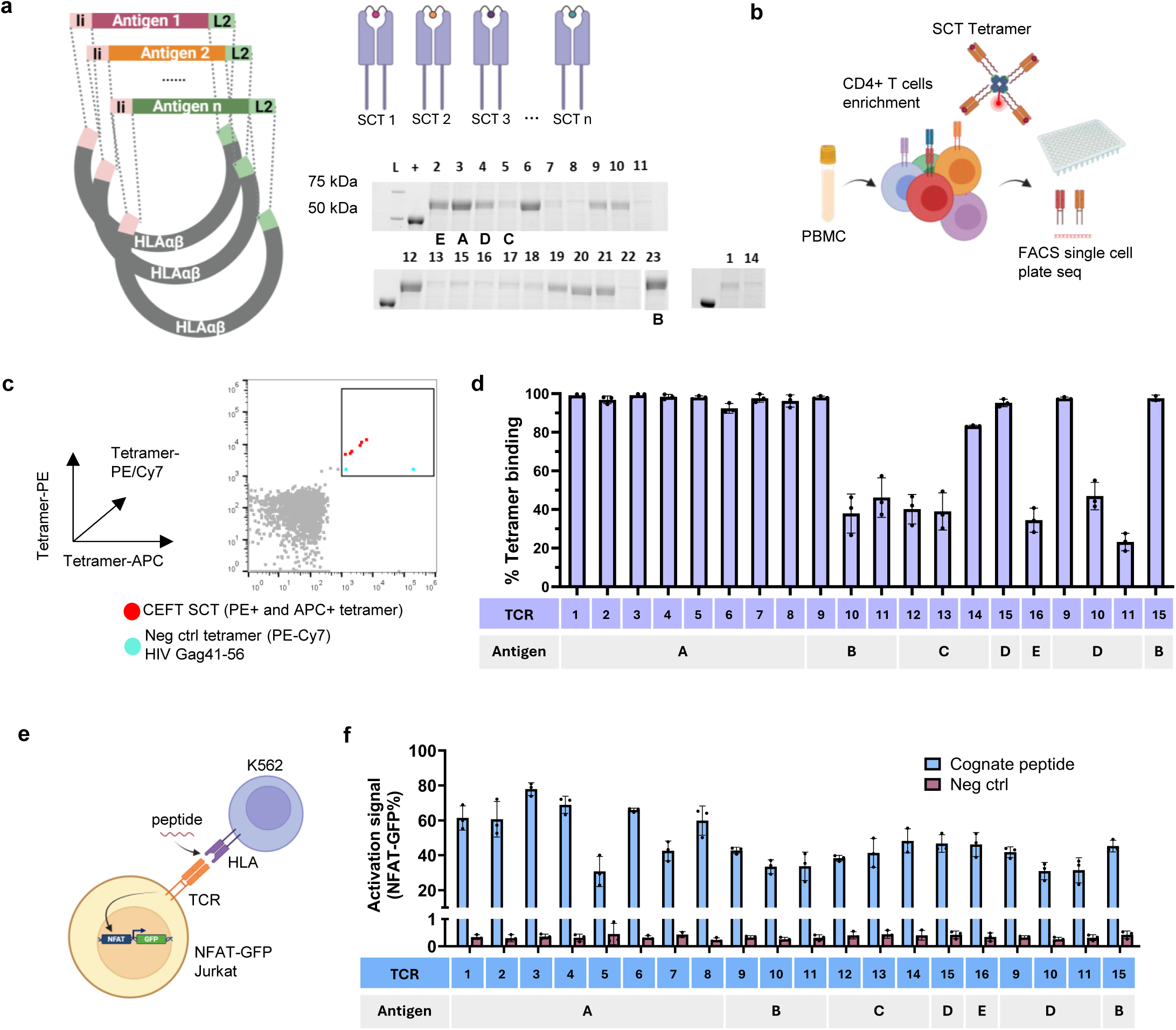
SCT library-based approach for antigen-specific CD4+ T cell discovery in healthy donors. **a.** A 23-Element DRB1*01:01 CEFT SCT library was used to identify novel antigen-specific CD4+ T cells. SCT expression was evaluated on SDS-PAGE protein gels. L, molecular weight ladder; +, protein standard for quantification of expression level. **b.** Schematic workflow for using the CEFT SCT library to identify novel antigen-specific CD4+ T cells from healthy donor PBMCs. **c.** Antigen-specific T cells were detected through dual tetramer staining to enhance the true positive rate. An HIV Gag_41-56_ SCT tetramer was used to exclude non-specific binding cells. A representative flow cytometry plot is shown. **d.** Tetramer binding validation of 16 newly identified CD4 TCRs, each reactive to one of the five CEFT antigens (A-E) (*n=3*). Tetramer binding signals were background-corrected using negative control tetramer staining. Error bars represent standard deviation. **e.** Schematic of peptide-pulsed activation assay. TCR-transduced NFAT-GFP Jurkat cells were co-cultured overnight with peptides and DR1-K562 cells. **f.** T cell activation measured through the percentage of GFP+ Jurkats (*n=3*). *P* < 0.01 for each group relative to the negative control peptide, determined by one-tailed independent t-test assuming equal variances.

To validate that the direct *ex vivo* SCT library screen accurately identifies true antigen–TCR pairs, newly discovered TCRs were cloned into TCR^KO^ NFAT-GFP Jurkat cells (**Table S3**) and evaluated for SCT binding (**Fig. 2d**). We confirmed TCR recognition to cognate antigens, with varying binding intensities suggesting differential recognition capacities among TCRs. To further assess whether SCTs faithfully model antigen presentation on MHCII and pMHC interaction to TCR, we tested these TCRs in a peptide-pulsed activation assay (**Fig. 2e**) using DRB1*01:01-expressing K562 cells with an empty peptide-binding pocket (DR1-K562)^32,33,34^. Co-culture with cognate peptides induced NFAT-based activation in all TCR+ Jurkat clones (**Fig. 2f**). Clones 9, 10, 11 and 15 showed consistent tetramer binding and activation to peptides (B and D) that share the same core sequence (**Table S4**). Together, these results demonstrate that SCT libraries enable direct *ex vivo* discovery of novel, functionally validated CD4+ TCR-antigen pairs with high specificity and sensitivity.

### High-throughput screening of the entire SARS-CoV-2 receptor-binding domain identifies large-scale antigen-specific CD4+ T cells

We next harnessed class II SCT libraries for high-throughput, systematic screening of the full SARS-CoV-2 receptor-binding domain (RBD), enabling comprehensive identification and characterization of antigen-specific CD4+ T cell responses in a longitudinal patient cohort. Prior studies have demonstrated the RBD to be a dominant target of neutralizing antibodies^35,36,37,38,39^. Given the importance of CD4+ T cells in B cell maturation, we thus constructed a 54-element SCT library spanning the entire RBD in 4-amino acid increments (**Fig. 3a**) to systematically profile CD4+ T cell responses. Of these, 46 SCTs were successfully expressed in usable quantities (**Extended Data Fig. 2a, Table S5**). An additional 18 SCTs representing reported non-RBD epitopes from spike (S), membrane (M), and nucleocapsid (N) proteins were also included^14,40,41,42^ (**Table S5, Extended Data Fig. 2a**).

**Figure 3.**
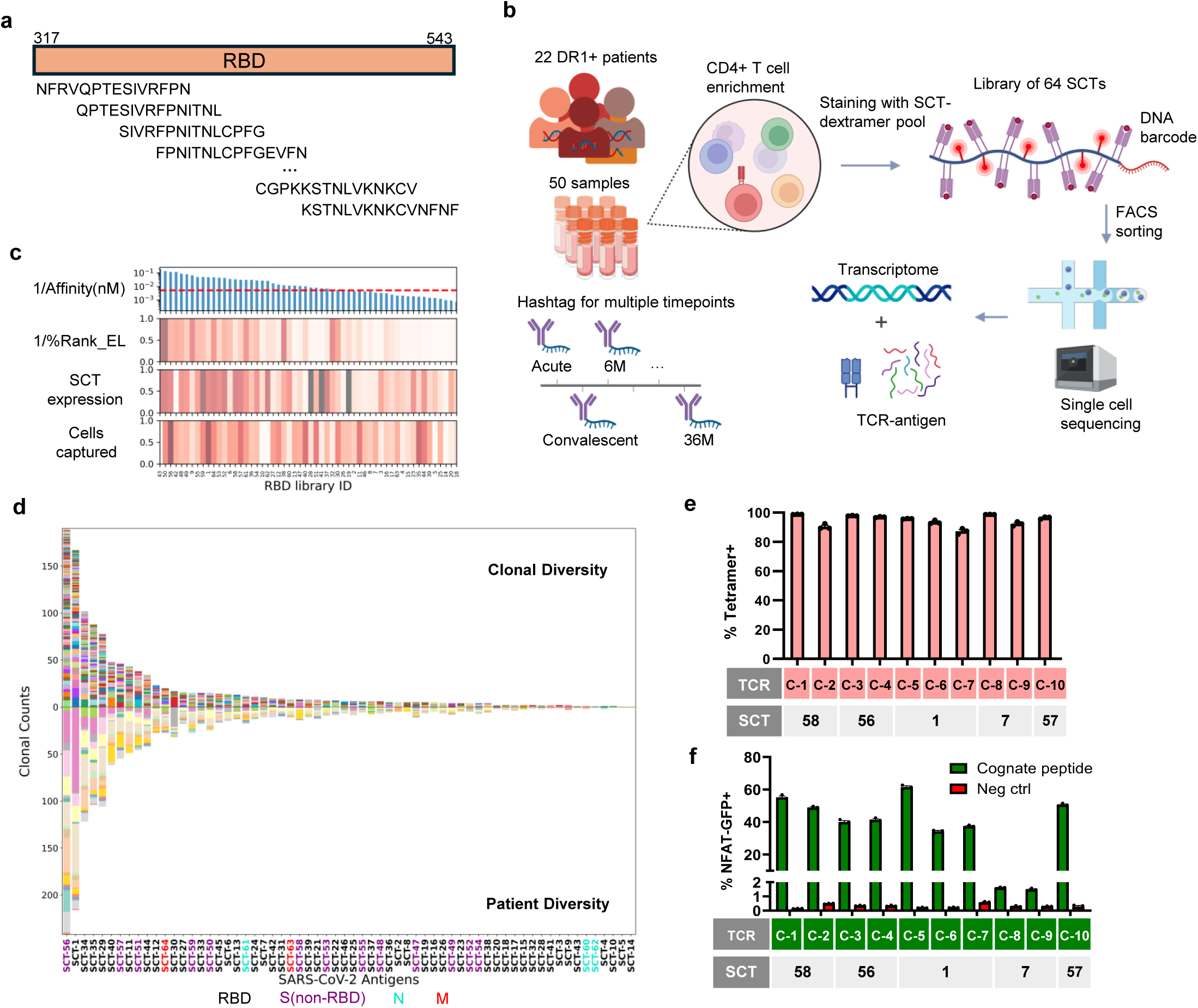
High-throughput screening of the entire receptor-binding domain (RBD) of SARS-CoV-2 spike identifies large-scale antigen-specific CD4+ T cells. **a.** Design of an SCT library with overlapping 15-mer peptides spanning the entire RBD of SARS-CoV-2 spike protein. **b.** Experimental workflow for high-throughput screening. PBMC samples from 22 HLA-DR1+ participants from the INCOV and UNCOVR cohorts were barcoded by timepoint, pooled, enriched for CD4+ T cells, stained with the 64-element SCT-dextramer pool, FACS sorted, and sequenced. Single-cell analysis included RNA expression profiling, TCRαβ sequencing, antigen-MHC pairing, and timepoint identification. Comparison of genetic variants between transcriptome and whole genome sequencing (WGS) was used to demultiplex patient identity (Methods). **c.** Correlation between SCT expression levels, the number of captured antigen-specific cells, and predictions from class II antigen-presentation algorithms. %Rank_EL, percentile rank of eluted ligands. SCT expression was quantified prior to purification. **d.** Clonal and patient diversity in responses to each SARS-CoV-2 antigen. Antigens are color coded based on their parent protein. **e.** Tetramer binding validation of SARS-CoV-2-specific CD4 TCRs (*n=3*). **f.** Peptide-pulsed activation of SARS-CoV-2 CD4 TCRs in NFAT-GFP Jurkats *(n=3). P* < 0.0001 for each group relative to the negative control peptide, determined by one-tailed independent t-test assuming equal variances.

The complete 64-member SCT library was multimerized using fluorophore-labeled, DNA-barcoded dextramers and pooled to stain CD4+ T cells from 50 PBMC samples across 22 DRB1*01:01-positive participants (**Fig. 3b**). Samples spanned multiple time points, ranging from acute infection (AC, <1 week of infection) through convalescence (CV, 2-3 months post-acute) to long-term follow-up (6 to 36 months). Sample multiplexing was enabled by hashtag antibodies, facilitating time-resolved deconvolution. Dextramer-positive CD4+ T cells were isolated by FACS and subjected to single-cell sequencing, yielding paired whole transcriptome, TCR sequence, epitope specificity, HLA restriction, and patient/timepoint origin (**Fig. 3b**). After data processing (**Methods**), we identified a total of 2,188 antigen-specific CD4+ T cells, with cells detected across all donors.

Analysis of SCT expression revealed limited correlations with in silico MHC-binding predictions (**Fig. 3c, Extended Data Fig. 2b**), consistent with known challenges in prediction algorithms. Similarly, cell capture was independent of SCT expression level, predicted binding affinity, or percentile rank, in contrast to class I SCT platforms^43^. These results underscore a key advantage of class II SCT libraries for empirical screening.

The broad diversity of T cell clonotypes captured by each SCT, as well as the distributions of antigen-specific CD4+ T cell across participants, confirmed that the observed responses were not skewed by donor, clonotype, or antigen bias (**Fig. 3d**). Clonal expansion was observed in responses to 42 of 64 antigens and in 49-58% of cells across time points (**Extended Data Fig. 2c**). We further validated epitope-specific TCRs through tetramer binding and peptide-pulsed activation on selected clonotypes (**Fig. 3e and 3f, Table S6**), confirming the fidelity of the SCT-based high-throughput identification strategy.

### Multi-modal and high-dimensional profiling empowers deep characterization of CD4+ T cells at single-cell resolution

The SCT-library approach permits comprehensive characterization of thousands of antigen-specific CD4+ T cells by integrating single-cell resolution data across multiple dimensions (**Fig. 4a, Methods**). Each T cell is deeply profiled through three parallel sequencing libraries: 1) whole transcriptome, 2) TCR αβ sequences, and 3) a surface protein library which decodes MHC-restricted antigen specificity and timepoint origin via DNA-barcoded dextramers and hashtag antibodies. Patient identity is resolved by matching transcriptome-derived genetic variants to corresponding whole-genome sequencing profiles.

**Figure 4.**
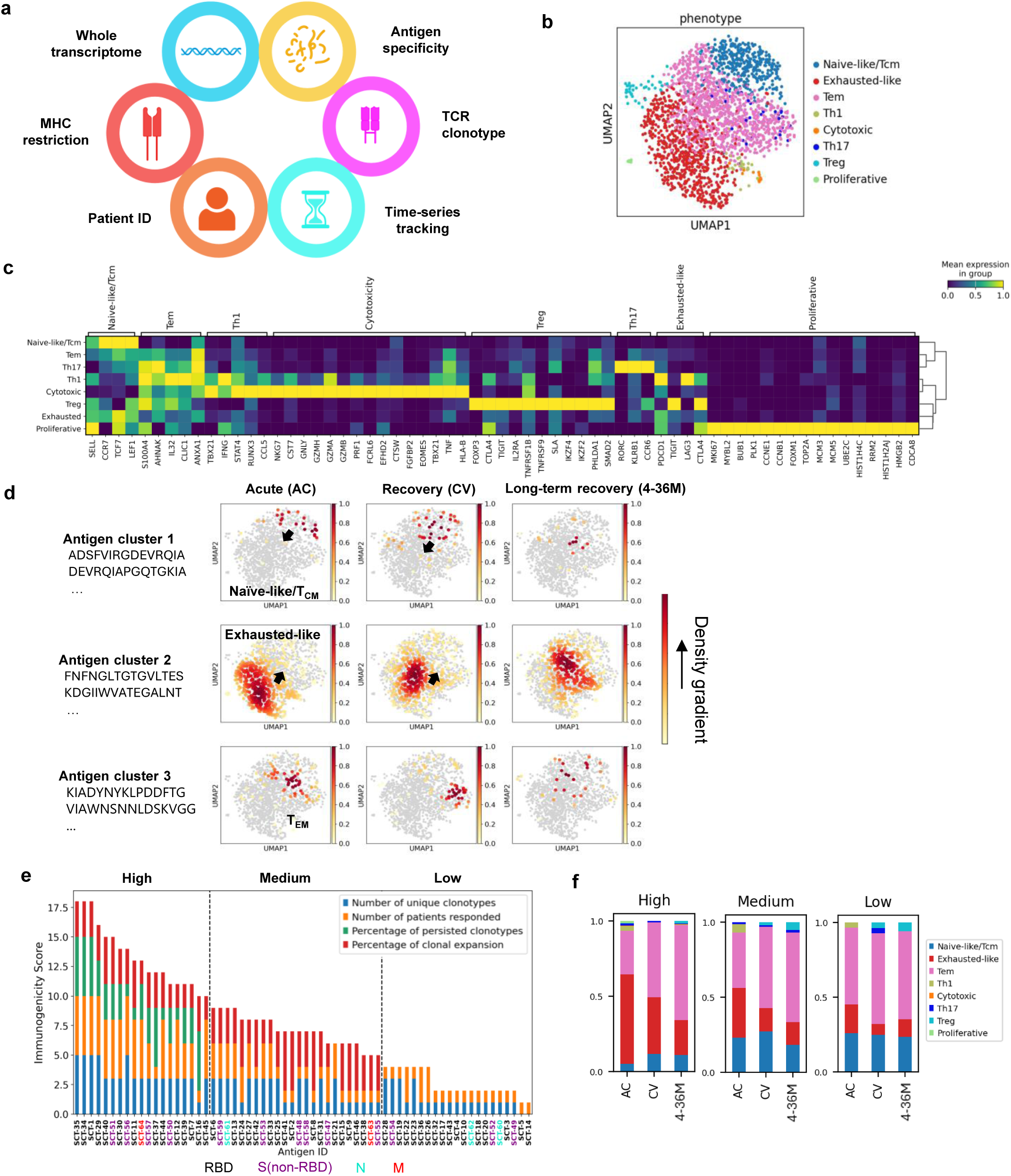
Multi-modal, high-dimensional profiling empowers deep characterization of CD4+ T cells at single-cell resolution. **a.** Graphical description of the high-dimensional data extracted from each single antigen-specific CD4+ T cell. **b.** UMAP projection of single cell transcriptomic profiles from SARS-CoV-2-specific CD4+ T cells. **c.** Expression of signature gene markers used to annotate Leiden-defined clusters. Expression levels were normalized across clusters. **d.** Longitudinal trajectories of phenotypic states of antigen-specific CD4+ T cells across timepoints. **e.** Immunogenicity ranking of DRB1*01:01-restricted SARS-CoV-2 antigens. Each property contributing to immunogenicity was scored on a log2 scale (0 to 5) and summed to generate a composite score. **f.** Phenotypic evolution of CD4+ T cells induced by antigens of low, medium, and high immunogenicity.

To dissect CD4+ T cell states, all antigen-specific CD4+ T cells were clustered by gene expression similarity and projected onto a Uniform Manifold Approximation and Projection (UMAP) (**Fig. 4b, Methods**). Phenotypic annotation based on canonical markers identified major subsets including naïve-like/central memory T_CM_ (SELL, CCR7, TCF7, LEF1)^44,45,46,47^, effector memory T_EM_ (S100A4, AHNAK, IL32, and CLIC1)^45,48,49^, Th1 (TBX21, IFNG, STAT4, RUNX3, CCL5)^1,45^, Treg (FOXP3)^1,44,45^, and Th17 (RORC, KLRB1, CCR6)^45^ (**Fig. 3c**). A subset of Th1 cells upregulates cytotoxic markers (NKG7, CST7, GNLY, PRF1, TNF, granzymes)^45,46,50^, while the exhausted-like phenotype is characterized by high expression of exhaustion markers (PDCD1, TIGIT, LAG3, CTLA4)^46^. Additionally, a small cluster of cells displays high expression of proliferation markers, including MKI67, MYBL2, BUB1, PLK1, and CCNE1^51^. These clusters reveal distinct phenotypic variations among SARS-CoV-2-specific CD4+ T cells.

A three-dimensional characterization of CD4+ T cells using epitope, timepoint and transcriptional information revealed important biological phenomena. Antigen-induced responses were phenotypically polarized at early timepoints, with distinct biases toward naïve-like/T_CM_, exhausted-like, and T_EM_ (**Fig. 4d, Methods**). Longitudinal tracking revealed a convergence toward T_EM_ phenotypes, consistent with previous reports on the evolution of CD4+ T cell responses following antigen stimulation^52,53^.

To showcase the full potential of this powerful dataset for fundamental immunology studies, we conducted an integrative analysis incorporating five key aspects—MHC and epitope specificity, TCR clonotype, patient identify, and timepoint origin—enabling a novel investigation into the immunogenicity of class II–restricted antigens. We established a quantitative scoring framework incorporating: (1) number of unique TCR clonotypes per antigen, (2) donor response frequency, (3) longitudinal persistence, and (4) clonal expansion (**Methods**). These metrics were log_2_-transformed, aggregated, and used to rank antigens into high, medium, and low immunogenicity categories (**Fig. 4e, Extended Data Fig. 3a**). Low to moderate inter-parameter correlations (ρ = 0.23-0.64) indicated that each parameter retains a degree of independent information (**Extended Data Fig. 3b**).

By folding in the sixth key parameter—transcriptomic information—we found that highly immunogenic antigens elicited more exhausted-like CD4+ T cell responses during acute disease, followed by the most pronounced phenotypic transition to effector memory over time. In contrast, low-immunogenicity antigens induced naïve-biased responses with limited evolution (**Fig. 4f**). Overall, the platform inspires a scalable, data-driven framework for assessing class II antigen immunogenicity, potentially aiding future vaccine designs for more targeted immune responses.

A further review of clonal architecture revealed that Th1 and proliferative subsets were enriched for expanded clonotypes, while naïve-like cells remained largely unexpanded (**Extended Data Fig. 3c**). Persistent clonotypes were enriched in exhausted, TEM, cytotoxic, and Th17 compartments. Public TCRs were detected across individuals. In one case, donor INCOV042, longitudinal sampling revealed novel clonotypes emerging at late timepoints, coinciding with persistent symptoms, in line with recent reports of viral antigen persistence in long-COVID symptoms^54,55,56^ (**Extended Data Fig. 2d**).

Collectively, this high-throughput, multi-modal approach enables systematic identification and deep characterization of class II-restricted antigen-specific CD4⁺ T cells at single-cell resolution, revealing critical immunological mechanisms that drive disease immunity.

### Whole-proteome screening of HPV-16 E6/E7 and TCR repertoire profiling uncover diverse CD4 TCR functionality in precancerous patients

Next, we explored the application of class II SCT-based platform in cancer, performing proteome-wide screening of the oncogenic HPV-16 proteins E6 and E7 for CD4 TCR repertoire profiling. HPV-16 is a high-risk genotype and the primary drive of cervical, oropharyngeal, anal, and vaginal cancers. While HPV-specific CD8+ TCR-T therapies have demonstrated clinical benefit^57,58,59,60^, the contribution of HPV-reactive CD4+ T cells remains less well characterized, despite growing evidence of the critical role of CD4+ T cells in anti-tumor immunity^3^. E6 and E7 proteins are universally expressed in HPV-16-transformed cells, and hence, ideal targets for CD4 TCR repertoire profiling and providing mechanistic insight into class II antigen recognition.

A major challenge for a comprehensive whole-proteome screening lies in the complexity of antigen presentation on class II MHC. Unlike class I MHC, which typically present peptides of fixed length, class II MHC has an open binding pocket that accommodates variable-length antigens with a 9-mer core flanked by peptide flanking regions (PFRs). Recent studies suggest that PFRs may modulate CD4+ TCR recognition, yet their functional relevance remains poorly understood^61,62,63,64^. To systematically investigate this, we designed a comprehensive 92-element SCT library spanning the full E6 and E7 proteome, covering all possible lengths (13-25-mer) of putative antigens, with special focus on screening peptide families of varying lengths that share the same core antigen (**Fig. 5a, Methods, Table S7**). Of these, 85 SCTs reached expression levels sufficient for purification and downstream screening.

**Figure 5.**
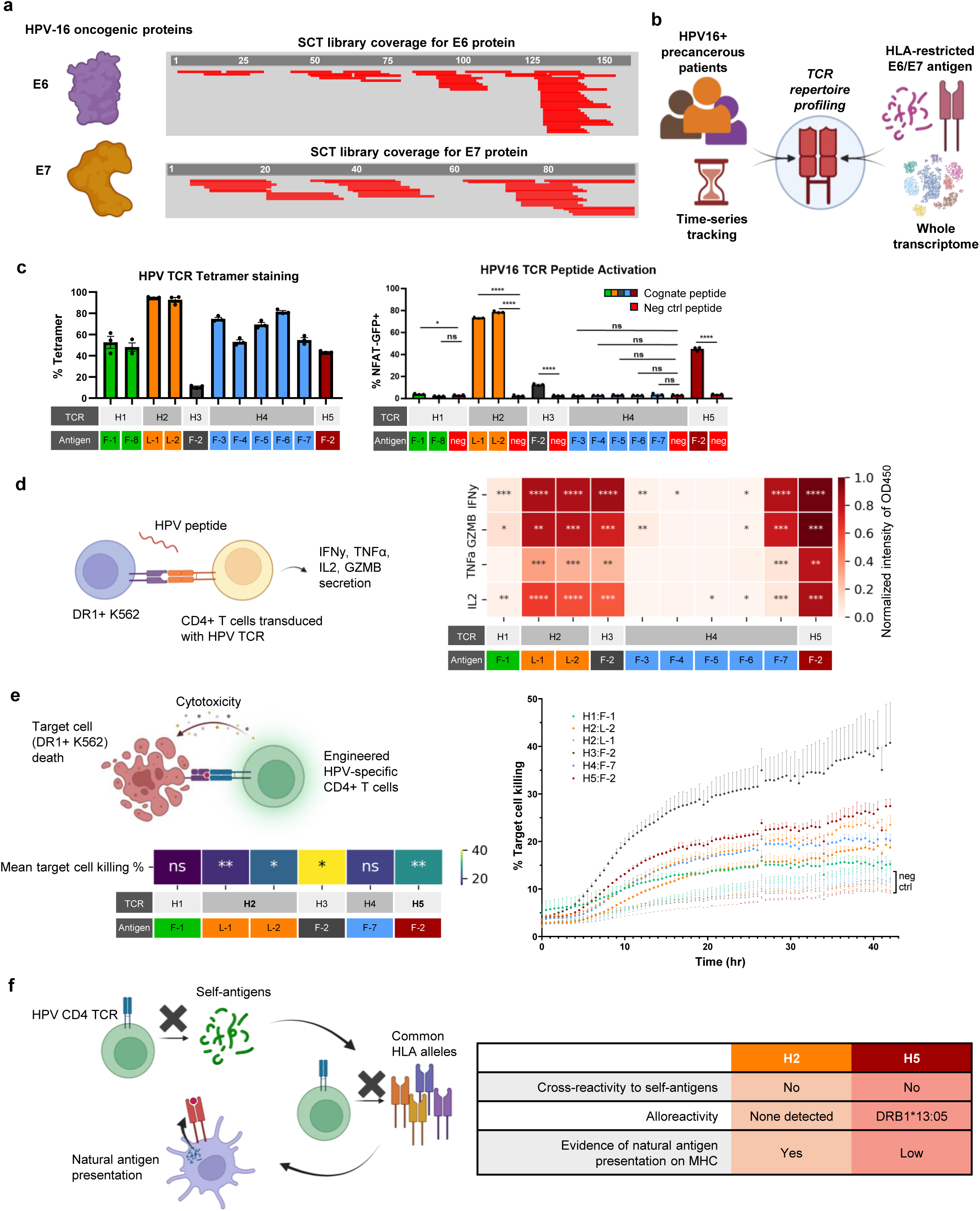
Whole-proteome screening of HPV-16 E6 and E7 enables rapid TCR repertoire profiling and comprehensive mapping of TCR functional heterogeneity. **a.** Antigen coverage across the E6 and E7 proteins in the SCT library for proteome-wide screening. **b.** Schematic overview of TCR repertoire profiling integrated with other key modalities. **c.** Validation of HPV-specific CD4 TCRs (H1-H5) through tetramer binding and peptide-pulsed activation assays *(n=3).* \*\*\*\**P* < 0.0001, \**P* < 0.05, ns *P* >0.05 labeled for each group relative to the negative control peptide, determined by one-tailed independent t-test assuming equal variances. **d.** Polyfunctionality TCR-transduced primary CD4+ T cells assessed by cytokine production (IFNγ, TNFα, IL2, and GZMB) via ELISA assay following antigen stimulation (*n=3*). OD_450_ values was normalized for each cytokine. \*\*\*\**P* < 0.0001, \*\*\**P* < 0.001, \*\**P* < 0.01, \**P* < 0.05, and unlabeled tiles indicate *P* ≥ 0.05, all relative to negative control peptide. **e.** Cytotoxicity of TCR-transduced CD4+ T cells against DR1-K562 cells pulsed with cognate HPV-16 E6 antigens, monitored over 42 hours *(n=2*). Peptide-mismatched controls were used for each TCR as negative controls. **f.** Comprehensive functional profiling of HPV-specific CD4 TCRs assessing cross-reactivity, alloreactivity, and evidence of natural antigen presentation on MHC.

We applied this library to PBMCs from a longitudinal cohort of patients with prolonged HPV-16 infection and high-grade cervical intraepithelial neoplasia (CIN II/III), including participants from an HPV-16 vaccine trial (n=3, **Extended Data Fig. 4a**) and an artesunate treatment study (n=1)^65^. PBMCs were prelabeled with hashtag antibodies for patient and timepoint demultiplexing, stained with DNA-barcoded SCT dextramers, sorted, and subjected to single-cell sequencing to capture TCR repertoire, transcriptome, and antigen specificity (**Figure 5b, Methods**). UMAP clustering annotated distinct CD4+ T cell phenotypes (**Extended Data Fig. 4b, 4d**). Notably, the exhausted-like phenotype, which was prominent (and known^46^) in the SARS-CoV-2 specific CD4+ T cells (**Fig. 4b**) were absent in this context.

Across the cohort, we identified 112 HPV-16 specific CD4+ T cells. We selected a subset of 25 CD4 TCRs for in-depth functional characterization, with a preference for clonally expanded and persistent clonotypes, and identified five CD4+ TCRs (H1-H5) as E6-specific (**Extended Data Fig. 4c, Table S8**). We first validated their specificity through tetramer binding (**Fig. 5c, left**). TCRs H1, H2, and H4 each recognized multiple peptides sharing the same 9-mer core (**Table S8**), underscoring the role of the core epitope in TCR engagement^66^. Notably, H1 and H3-H5 are TCR clonotypes that recognize variants of a peptide family sharing the same core. Surprisingly, peptide-pulsed activation of TCR-transduced NFAT-GFP Jurkats revealed that tetramer binding does not always translate into downstream activation: H1 and H4 bound SCTs robustly but failed to trigger downstream signaling (**Fig. 5c, right**), highlighting the potentially critical role of PFRs in modulating functional outcomes.

To further investigate the asscoiation between PFRs and T cell functional capacity, we transduced these TCRs into TCR^KO^ primary CD4+ T cells enriched from healthy donors and confirmed their specificity via tetramer staining (**Extended Data Fig. 4e, 4f**). Notably, the knock-in efficiency was markedly lower in CD8+ than CD4+ T cells (**Extended Data Fig. 4g**). We then evaluated these TCRs through a comprehensive series of functional assays. We first assessed their cytokine secretion in response to peptide-pulsed DR1+ K562 cells (**Fig. 5d**). TCRs H2, H3, and H5 displayed on-target secretions of all cytokines, while H4 showed minimal secretion for most peptides except F-7, despite strong tetramer binding to the broader peptide family sharing the same core (F-3 to F-7). These results further emphasize the influence of PFRs on CD4+ T cell activity.

Given emerging evidence of cytotoxic CD4+ T cells in anti-tumor immunity^3^, we assessed the cytotoxicity of HPV-specific TCRs. TCR-transduced CD4+ T cells were co-cultured with DR1+ K562 cells pulsed with cogante peptides and labeled with Cytolight Rapid Dye (**Fig. 5e**). H2, H3, and H5 induced significant target cell apoptosis, wherea as H1 and H4 showed minimal killing.

Based on functional performance, TCRs H2 and H5 were pritorized for further pre-clinical evaluation (**Figure 5f**). To assess off-target effects, we performed alanine scanning of their 9-mer core epitopes, identifying key recognition motifs: *xxxxYNKPx* (H2) and *FHNxRGRWx* (H5) (**Extended Data Fig. 4h**). BLAST-based homology search identified human self-antigens with similar motifs (**Table S9**). Co-culture assays using HLA-matched, EBV-transformed B-lymphoblastic cell lines (LCLs) pulsed with these peptides showed no evidence of cross-reactivity, indicating strong target specificity (**Figure 5f, Extended Data Fig. 4i**).

We then screened for alloreactivity to common class II HLAs using a panel of 41 LCLs expressing 92 distinct class II HLA alleles. H5 exhibited reactivity toward two LCL lines sharing the HLA-DRB1*13:05 allele, whereas H2 demonstrated no evidence of alloreactivity (**Fig. 5f, Extended Data Fig. 4j, Table S10**). Finally, titration of the full-length E6 protein pulsed to a DRB1+ LCL line showed functional activation of H2-transduced T cells, confirming that the L-1 antigen is naturally processed and presented by DRB1*01:01 (**Fig. 5f, Extended Data Fig. 4k)**. Thus, the H2 TCR emerges with strong therapeutic potential for clinical translation.

Together, these results demonstrate the power of class II SCT libraries for proteome-scale screening of functional CD4+ TCR repertoires in cancer. This approach enables precise mapping of peptide families, functional interrogation of PFRs, and identification of therapeutically relevant CD4+ TCRs with minimal off-target risk—laying the groundwork for translational CD4⁺ T cell immunotherapies.

## Discussion

CD4+ T cells are essential orchestrators of adaptive immunity, with critical roles in infectious disease, cancer, and autoimmunity. However, systematic and deep characterization of antigen-specific CD4+ T cell responses at scale remains technically challenging. Conventional approaches either have limited throughput or lack resolution of key biological parameters, including MHC restriction, phenotypic state, and patient/timepoint identity.

We reported here on a scalable platform using class II SCT libraries to enable high-throughput, whole-proteome screening for antigen-specific CD4+ T cells with associated co-profiling of epitope specificity, MHC restriction, TCRα/β genes, whole transcriptome, patient identity, and time-series tracking. We benchmarked the performance of individual SCTs, showing a comparable specificity and sensitivity to conventional pMHCs. We then demonstrated the validity of SCT library-based approach to discover novel antigen-CD4 TCR pairs in healthy donor PBMCs. To demonstrate the power of the platform in high-throughput multi-modal profiling, we performed full-length screening of the SARS-CoV-2 spike receptor-binding domain in a longitudinal cohort of COVID-19 participants. The resulting novel, high-dimensional dataset permitted comprehensive kinetic analysis of antigen-specific CD4+ T cells, and informed a multi-component metric for defining class II-restricted immunodominance. Given the importance of CD4+ T cells in B cell maturation, the ability to quantitate immunogenic viral antigens should provide a compelling approach towards enabling rational and precision vaccine design. Notably, the five most immunogenic antigens identified in this study are all RBD-derived epitopes—a trend consistent with previous findings highlighting the critical role of anti-RBD IgGs in conferring resistance to SARS-CoV-2^36^.

We further tested the versatility of the platform in CD4 TCR repertoire profiling in cancer settings. We performed a proteome-wide screen for HPV-16 E6 and E7 specific TCRs in patients with pre-cancerous HPV+ lesions and identified a panel of HPV-specific CD4+ T cells. Extensive functional analysis revealed peptide-flanking region–dependent effects on TCR binding, activation, cytokine production, and cytotoxic function. These findings underscore the importance of comprehensive functional characterization for TCRs targeting class II–restricted antigens in immunotherapeutic development.

A current limitation of our work is the focus on a single class II allele for SCT libraries, DRB1*01:01 We have demonstrated that SCTs can be reliably constructed and expressed across multiple common alleles, including DPB1*04:01, DRB1*07:01, and DRB1*11:01. Expanding SCT libraries to these alleles would significantly increase population coverage but may require template optimization for certain alleles. Such work is currently underway. As SCT-based datasets accumulate, they may ultimately provide high-quality training data to advance in silico models for predicting CD4+ T cell antigen recognition.

Collectively, we demonstrated a versatile, high-throughput platform that empowers proteome-scale identification and multi-modal characterization of antigen-specific CD4+ T cells. This method resolves key biological variables at single-cell resolution and offers a generalizable solution for dissecting antigen-specific CD4+ T cell responses in diverse contexts, including infectious diseases, tumor immunology, autoimmunity, vaccine design, and fundamental T cell immunobiology.

## Acknowledgements

We would like to thank all the participants in the InCoV and UNCOVR studies and the HPV clinical trials for their support and contributions. We thank Y. Su for his advice and guidance on project planning and manuscript revision; P. Troisch for her support in preparing sequencing libraries and sequencing runs; the ISB Biobank team for collecting and processing clinical samples; the Flow Core and team members at Fred Hutchinson Cancer Research Center and Benaroya Research Institute. We thank M. van Meurs, N. Gu, and K. Wang for their assistance in the projects; F. Gao, V. Adit, and M. Davis for generously sharing plasmid sequences, reagents, and advice on troubleshooting protocols. This work was supported by the Andy Hill Cancer Research Endowment (CARE) Fund under award BRK_201801-05 and the NIH R01 grants (CA264090-01 to J.R.H. and CA263256 to C.L.T.). K.G.P. was supported by the Paul G Allen Research Center at Providence-Swedish Cancer Institute. Y.S. was supported by the Damon Runyon Quantitative Biology Fellowship from the Damon Runyon Cancer Research Foundation (DRQ-13-22).

## Author Contributions

Conceptualization and design, R.Z., W.C., Y.S., and J.R.H.; Acquisition, analysis, or interpretation of data, R.Z., J.Q., M.M., J.C., V.G., C.B., S.H., J.X., D.Y., A.W., S.K.S., A.A., D.C., Y.R., Y.S., and J.R.H.; Creation of new software used in the work, R.Z., W.C., R.N., J.X., D.C., and S.W.; Manuscript writing and revision, R.Z., Y.S., and J.R.H.; Clinical sample acquisition, R.E., K.M.M., S.L., C.M., L.R., J.R.W., J.C., K.N., J.F., A.D., S.J.F., M.S., S.P., C.E.B., A.R., D.W., K.G.P., C.W.D., C.P-S., J.D.G., C.L.T., .and J.R.H.

## Methods

### Ethical statement

The research presented here complies with all relevant ethical regulations. Procedures for the INCOV study were approved by the Institutional Review Board (IRB) at Providence St. Joseph Health with IRB study number STUDY2020000175 and the Western Institutional Review Board (WIRB) with IRB study number 20170658. The samples from the HPV vaccine study were collected from a phase 2b trial registered at ClinicalTrials.gov (number NCT01304524) and EudraCT (number 2012-001334-33). The protocol was approved by the institutional review board or ethics committee at each participating center, and all patients gave written informed consent. An independent data and safety monitoring board reviewed unmasked safety and histopathological regression data.

### Primer auto-generation algorithm

The algorithm (https://github.com/rzhang101/classII_sct_primer_auto_generation.git) takes in a user-defined list of antigens in one-letter coded amino acid sequences. An initial gene fragment including the DNA sequences of the antigen generated through reverse translation and codon optimization, the pIi domain, and L2 are constructed. An initial set of forward and reverse primers are designed to allow at least 16 base pair (bp) of hybridization. The set of primers is then tested through three checkpoints to confirm PCR effectiveness: 1) if the melting temperature of the 20 bp hybridization region is between 50 to 69C using Nearest Neighbor thermodynamics; 2) if at least 16 bp are annealed at the annealing temperature and if the annealed primers are the majority by analyzing the primer structures through NUPACK^67^; 3) if the two primers have at least 16 bp annealed at the extension temperature and if a hairpin structure exists that might hinder the extension process. If any of the checkpoints fail, the primers undergo a random selection of alternative codons until a codon combination passes all checkpoints. The algorithm outputs a list of forward and reverse primers ready for purchase.

### Class II SCT expression

The plasmid used for the expression of class II SCTs was built based on a pcDNA3.1 vector (Invitrogen, V86020). An SCT plasmid template encoding for a class II allele was assembled by a two fragment Gibson assembly each encoding for the alpha or beta chain of the allele. The antigen fragment was produced by an overlap extension PCR of the forward and reverse primer designed by the primer auto-generation algorithm (IDT). The PCR was carried out in a 20 uL reaction containing 1.5 pmol of each primer, 7 uL nuclease free water, and 10 uL KOD Hot Start Master Mix (Millipore Sigma, 71842). The amplified fragment was purified (Qiagen, 28104) and Gibson (New England Biolabs (NEB), E2621S) assembled into the plasmid backbone containing the HLA allele in a 5:1 insert to vector ratio or 0.25 pmol per fragment and 0.05 pmol of vector. The Gibson product was then transformed into Top10 competent cells (Invitrogen, C404003) and the colonies were picked and cultured, followed by extraction of the amplified plasmids (27104). Plasmid sequences were confirmed by Genewiz, MCLAB, or Plasmidsaurus.

The mammalian cell based Expi293 transfection system (Gibco, A14635) was used for SCT expression. SCT transfection was performed in a 24-well plate format for high-throughput production. Briefly, for each SCT, 1.25 mL of Expi293 cells at 3M/mL were seeded and transfected with the SCT plasmid packaged based on the manufacturer’s protocol. After 18-24 hour of incubation, Enhancer 1 and 2 were added followed by an incubation of 72 hours. The secreted SCT proteins were collected four days after the initial transfection. An aliquot of the supernatant was denatured to quantify the SCT expression level through SDS-PAGE (Bio-Rad, 4568035). Relative intensity of the SCT expression subtracting negative control was recorded. Supernatant of expressed SCTs were concentrated and buffer-exchanged with 20 mM bicine (pH 8.3) (Bufferad, B2239-83), and then biotinylated for 1.5 hour at room temperature and then overnight (Avidity, BirA500). The SCTs were his-tag purified on the next day (Biotage, PTR-92-20-03) and de-salted into PBS (Thermo Fisher, 89883). Concentration of the SCT protein was measured through nanodrop and stored in 20% glycerol/PBS. The CEFT SCT library was designed based on the antigen list used for ELISpot stimulation on healthy donor PBMC samples from ImmunoSpot.

### N-linked de-glycosylation assay

SCTs were de-glycosylated according to manufacturer’s protocol (NEB, P0704). Specifically, 2 ug of each purified SCT was incubated at 100C for 10 minutes with the 10X denaturing buffer, and chilled on ice. PNGase enzyme, 10% NP-40, GlycoBuffer, and water were mixed well and incubated at 37C for one hour. The PNGase-processed proteins were then reduced and run on a SDS-PAGE protein gel in comparison with non-PNGase processed SCTs.

### Cell line and media

Wild-type Jurkat E6-1 was purchased from ATCC (TIB-152). TCRb followed by TCRa knock-out was performed using CRISPR gRNA (IDT) and nucleofection, confirmed through flow cytometry, and sorted for purity. A plasmid encoding for NFAT-GFP reporter (VectorBuilder) was then lentivirally transduced into the TCR-Jurkats and treated with blasticidin (InvivoGen, ant-bl-05) for selection. DR1+ K562 cells were established from WT K-562 (ATCC, CCL-243), which lack functional expression of wild-type class II pMHC^68^, by lentivirally transducing a plasmid encoding for the DRA/DRB1*01:01 allele with the transmembrane and cytoplasmic tail domains. Expression of DR1 on the surface was confirmed through flow cytometry and FACS sorted for purity. HEK293T cells were purchased from ATCC (CRL-3216). Healthy donor PBMC samples were purchased from ImmunoSpot.

### TCR cloning

Fragments encoding for the V(D)JC regions of alpha and beta chains were designed, codon-optimized and purchased from Twist Bioscience. The fragments were PCR amplified and Gibson assembled into a pRRL plasmid (generously provided by Dr. Philip Greenberg, Fred Hutchinson Cancer Center) for lentiviral production. The sequence-verified plasmid DNA was transfected into the HEK293T cell line along with packaging plasmids to produce lentiviral particles (Qiagen, 301425) which were transduced into the target cell line. 48-72 hours after removal of the virus soup, the TCR expression was measured through flow cytometry.

### Tetramer binding assay

Tetramers were prepared by incubating the SCTs with a fluorophore-labeled streptavidin (Invitrogen, SA10041, SA1005, SA1012) in a 4:1 ratio. Usually 2.5 pmol of SCT and 0.625 pmol of streptavidin was used to generate tetramers for one staining reaction. After a 30-minute incubation on ice, 1 uL of 20 uM biotin was added to the tetramers followed by an incubation of at least 30 min on ice. Cells were incubated in 50 nM Dasatinib (PKI) (Adooq, A10290) at 37C for 20-30 min and then centrifuged. Cells were then stained with tetramer at 4C for 15-25 min in 50nM PKI and washed, followed by live dye (Invitrogen, C34858) and TCR-APC (Biolegend, 306718) staining. Cells were then washed and analyzed through flow cytometry.

### Antigen-specific CD4 T cells staining

The DRB1*01:01 SCT with antigen derived from the HIV Gag protein (HIV Gag_41-56_ SALSEGATPQDLNTML) was used as the negative control for SCT tetramer staining. SE buffer for cell and tetramer staining was prepared by adding 0.5% BSA and 2 mM EDTA and filtered. PBMCs were thawed in R10 media and pre-enriched for CD4+ T cells using magnetic-activated cell sorting (MACS) according to the manufacturer’s protocol (Miltenyi, 130-096-533). In the final step of CD4+ T cell enrichment, cells were centrifuged and resuspended in 1 mL of media. The enriched CD4+ T cells were then incubated in 50 nM PKI at 37°C for 20 minutes, followed by centrifugation at 500 g for 5 minutes. Staining solution for each antigen was prepared by adding 2 µL of PE and APC tetramers, along with 2 µL of HIV PE/Cy7 tetramer, to 100 µL of 50 nM PKI/SE buffer. Each sample was resuspended in 100 µL of the tetramer staining solution, incubated at 4°C for 20 minutes, washed and centrifuged. A cell surface staining buffer was prepared by adding 1 uL of CD4-BV421 (Biolegend, 317434) per 100 µL of buffer per 50k CD4+ T cells, and Calcein Green (Invitrogen, C3099) live stain at a final concentration of 0.2 µM. The cells were incubated for 10 minutes at 4°C. After staining, the cells were washed with 100 µL of SE buffer to remove excess reagents, centrifuged, and resuspended in 200 µL of PBS.

### Single-cell plate sequencing for TCR sequences

The cells were single-cell sorted into 96-well plates containing 12 uL of 1x lysis buffer (Takara, 635013) with RNAse inhibitor (Promega, N2515). The plates were spun down immediately after sorting and flash frozen on dry ice. When ready for sequencing the TCRs, the lysate plate was thawed on ice for 3 minutes and centrifuged at 1,000 g for 1 minute. A master mix of one step RT-PCR reaction was prepared according to the manufacturer’s protocol (Qiagen, 210212). Briefly, the following reagents were mixed for one reaction: 3.2 uL 5x buffer, 0.64 uL dNTPs at 10 mM, 0.64 uL enzyme mix, 5.12 uL RNAse free water, 0.55 uL of a mixture of the alpha or beta variable primers (1.75 uM), and 0.48 uL of alpha or beta constant primer at 10 uM. Each lysate was split into two reactions for amplifying the alpha and beta sequences separately. Following RT-PCR, cleanup was performed using 0.8x SPRI beads (Beckman Coulter, A63881). A second PCR further amplified the TCR fragment and extended the fragment with overlap regions for the third PCR that added in the barcoding regions for multiplexed sequencing through Illumina.

### Peptide stimulation assay

Peptides were purchased from GenScript and reconstituted in DMSO. 50k TCR+ NFAT-GFP Jurkat cells were co-cultured with DR1-K562 cells at a 1:1 ratio with 2 µg/mL peptides in 100 µL R10 media and incubated at 37°C for 16-24 hours. Cells were then stained with live cell dye Calcein UV (Invitrogen, C34858), TCR-APC (Biolegend, 306718), and HLA-DR-PE (Biolegend, 307605) and analyzed through flow cytometry.

### High throughput screening of antigen-specific CD4+ T cells

SCT-dextramer was prepared by mixing SCT and a DNA-barcoded dextramer-PE (ImmuDex) in a 22:1 molar ratio and incubated on ice for 30 min before adding 10 pmol of biotin for another 30 min incubation on ice. The dextramers were pooled together with negative control tetramer-PE-Cy7s. PBMC samples were stained with hashtag antibodies (Biolegend) based on patients and/or timepoints. After two washes, PBMCs were pooled for MACS-based CD4+ T cell enrichment through negative selection (Miltenyi, 130-096-533). Isolated CD4+ T cells were incubated in 50 nM PKI solution at 37C for 20-30 min, followed by dextramer pool staining in 50 nM PKI for 20-25 min at 4C. After 2-3 washes, the cells were stained with live dye (Biolegend, 427402), CD4-BV421 (Biolegend, 317434), and TCR-APC (Biolegend, 306718) for 10 min at 4C. The cells were then FACS sorted, washed and loaded onto Chromium Next GEM chips for single cell sequencing (10x Genomics). Gene expression, TCR, and surface protein libraries were prepared and sequenced on Illumina NextSeq2000.

### Single cell sequencing data analysis

The transcriptome, TCR sequences, surface protein levels, and antigen specificity were simultaneously assessed for each cell. Raw data were processed using the Cell Ranger Single-Cell Software Suite (v3.1.0, 10X Genomics) with GRCh38 as the reference genome. Gene expression data (GEX) were quality-controlled to eliminate low quality cells: cells with less than 200 or above 5,000 unique genes or a mitochondrial content above 15%. Any genes detected in less than three cells were also removed. Gene expression counts for each cell were normalized by the total expression, scaled by a factor of 10,000, and transformed to a log scale. All cells that pass the quality control were clustered through PCA and KNN graphs based on the highly variable genes and projected onto UMAPs. Phenotypes were defined based on enrichment of signature gene sets and assigned to single cells based on clusters identified using the Leiden algorithm. Single pair and orphan beta TCRs were selected and matched to the GEX data. In the SARS-CoV-2 study, patient identity was deconvoluted through comparison of single nucleotide polymorphisms (SNPs) in transcriptome data with WGS profiles of all patients (described in detail in the *Single cell demultiplexing using genetic variants* section). Any doublet cell was identified based on SNPs assignment to multiple patients and excluded from the analysis. DNA sequences encoded for hashtags were analyzed to determine the time origin of SARS-CoV-2 samples or both the time origin and the patient identify for the HPV samples. Raw reads for each hashtag and dextramer were normalized. The time origin and/or patient identify of a cell was assigned based on the hashtag with the highest expression, provided it exceeded the second-highest hashtag by at least 5%. Cells were assigned a specific antigen only if they had a dextramer UMI representing more than 20% of the cell’s total dextramer reads. Cells that could not be definitively assigned to a specific dextramer were discarded.

### Single cell demultiplexing of patient origin using genetic variants

Patient identity was resolved by comparing single nucleotide polymorphisms (SNPs) in transcriptome data with WGS patient profiles, using previously described bioinformatics pipeline^69^. In short, a multi-donor genotype reference of exon single nucleotide variants (SNV) was constructed from WGS data using GATK (v4.1.9.0)^70^. The same variants were determined for each single cell using via 10x Genomics VarTrix (https://github.com/10XGenomics/vartrix). Finally, vireo (v0.5.0)^71^ compares single cell SNVs with the donor genotype reference to assign each single cell to a specific donor or as a doublet.

### Phenotype evolution analysis

Antigens were clustered based on the phenotype distribution of their corresponding CD4+ T cells at the initial infection time point (AC). Antigens with fewer than three corresponding T cells were excluded from the analysis. For each antigen, the dominant phenotype of antigen-specific CD4+ T cells was identified and assigned as its initial phenotype cluster. Phenotype distribution of the T cell responses induced by the antigen clusters were analyzed by calculating the density of the cells using Gaussian kernel density estimation at each timepoint and projected onto a UMAP.

### Immunogenicity score analysis

Four unique properties were assessed for each antigen: (1) the number of donors reactive to the antigen, (2) the number of unique CD4 TCR clonotypes specific to the antigen (normalized for the number of donors), (3) the percentage of unique T cell clonotypes that persisted across at least two time points, and (4) the percentage of clonally expanded TCR clonotypes (adjusted for persistent and cross-donor TCRs). All properties were adjusted to minimize confounding influences. For each property, antigens were ranked from highest to lowest, divided into groups based on a log₂ scale, and assigned scores ranging from 0 to 5 (**Extended Data Fig. 3c**). The total scores for each antigen were mmed, and immunogenicity ranks (high, medium, or low) were assigned based on a log₂ scale of the overall score.

### HPV-16 E6 and E7 antigen selection

The class II SCT library encoding the E6 and E7 proteins encompasses 92 antigens varying from 13-25 amino acids. The antigens were selected based on the NetMHCIIpan4.0^72^ and MixMHC2pred^73^ predictions. The top 15% of the antigens predicted to bind the DRB1*01:01 allele were selected. Out of the 92 selected antigens, 85 SCTs were successfully expressed with a concentration applicable for downstream applications. The SCTs were constructed into a pool of dextramers where each DNA barcode corresponds to a unique SCT.

### HPV-TCR+ primary CD4 T cell construction

CD4+ T cells were extracted from healthy donor PBMCs via MACS sorting and resuspended at a density of 1 M/mL in Prime-XV media (Fujifilm Irvine Scientific, 91154) with 2% Physiologix serum replacement supplemented with IL-7 (Biolegend, 581904) and IL-15 (Biolegend, 570304), both at a final concentration of 12.5 ng/mL. Each 1M cells were activated by adding 10 µL of TransAct human CD3/CD28 activator (Miltenyi, 130-111-160,) and incubated for 48 hours. After activation, endogenous TCRs were knocked out through CRISPR-Cas9 gene editing and target TCRs were transduced into the primary CD4+ T cells through lentivirus.

### ELISA

DR1+ K562 cells were pre-loaded with peptide for a final concentration of 0.1 mg/mL and incubated at 37°C for 1-2 hours. Excess peptides were washed and removed. Peptide-loaded K562 cells were co-cultured with primary CD4+ T cells in a 1:1 effector-to-target (E:T) ratio for 16-20 hours. ELISA assays for IL2, INFγ, TNFα, and GZMB were performed according to manufacturer’s manual (R&D Systems, DY202-05, DY285B-05, DY210-05, DY2906-05).

### Cytotoxicity assay

The cytotoxic assay was conducted through IncuCyte. DR1-K562 cells were preloaded with target peptide in a similar way to the ELISA assays and seeded at 1M/mL. 10 µL of the 100X working solution of Cytolight Rapid Dye (Sartorius, 4705) was used for every 1M K562 cells. The cells were incubated for 20 minutes at 37°C, and gently mixed every 10 minutes to allow the dye to fully bind. Any excess dye was washed out and the supernatant was removed. The cells were then resuspended to a concentration of 500k cells/mL and 50k cells were seeded and allowed to settle at room temperature for 30 minutes. The apoptosis reagent was prepared by diluting Caspase-3/7 green (Sartorius, 4440) to a final concentration of 20 µM (4x) in culture media and added to cells. The immune effector cells (primary TCR-T cells) were added to the DR1-K562 cells at an E:T ratio of 1:1. The assay plate was placed in the IncuCyte Live-Cell Analysis System for a 44-hour repeat scanning.

### Alanine scanning

Peptides were generated for the E6_91-107_ epitope (TCR H2) and E6_129-142_ epitope (TCR H5) with each position replaced with an alanine amino acid (Genscript). Healthy donor T cells, expressing TCR H2, TCR H5, were co-cultured with a DRB1+ K562 cell line that were previously pulsed with a dose titration (0.001-1μM) of the respective peptides. After approximately 20 hours, supernatants were harvested from the co-cultures and measured for cytokine secretion (R&D Systems, DY285B-05).

### Cross-reactivity assessment

BLAST search and motif scan were conducted to identify human self-antigens containing motifs similar to those revealed by alanine scanning. Healthy donor T cells expressing the HPV-TCR of interest were then co-cultured with an HLA matched EBV-transformed B-lymphoblastic cell lines (LCLs) (Coriell Institute for Medical Research). For the TCR H2, single dose (1 µM) of each peptide was used. For the TCR H5, 1, 2, 10 µM of each peptide was used. After ∼20 h, supernatants were collected and IFNγ cytokine secretion was determined using the ELISA assay.

### Alloreactivity assay

TCR-transduced primary CD4+ T cells were co-cultured overnight with each of the 41 EBV-transformed B cell lines (LCLs) and assessed for IFNγ secretion. As positive controls, four DRB1*01:01+ LCL cell lines were pulsed with 10 uM of E6_91-107_ (TCR H2) or E6_129-142_ (TCR H5) peptides respectively, and IFNγ secretion was measured.

### E6 protein pulsed T cell functional activation assay

A DRB1*01:01+ LCL (GM17281B) was pulsed with E6 protein (Rndsystems, AP-120) at 10 μg/mL, 2 μg/mL, 1 μg/mL, and 0 μg/mL for 3 hours, and then co-cultured with H2-TCR-transduced CD4+ T cells overnight. IFNγ cytokine secretion in the supernatant was determined using the ELISA assay. A DRB1*01:01 negative LCL (GM12244A) was used as the negative control.

---

**Extended Data Figure 1.**
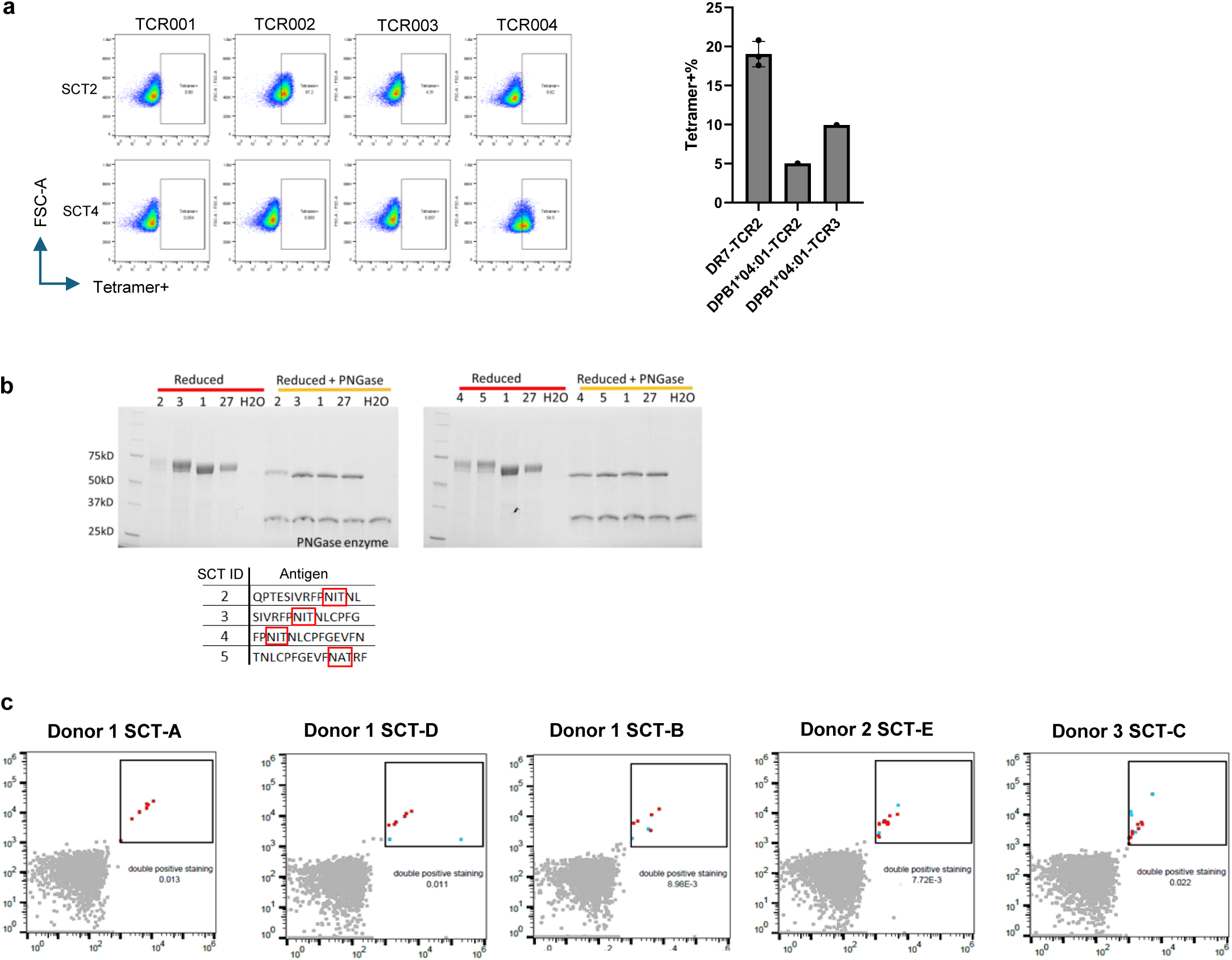
SCT construction and validation. **a.** Validation of class II HLA SCT templates using additional known antigen-TCR pairs. *left*, tetramer binding of DRB1*01:01 SCTs to matched TCRs. SCT2 paired with and TCR002 and SCT4 with TCR004. TCR001 and TCR003 served as irrelevant TCR controls. *right*, additional binding validation of DRB1*07:01 and DPB1*04:01 SCTs with their corresponding paired TCRs. **b.** De-glycosylation of N-linked glycosylation on SCTs. SCTs 2-5 contain NxT motifs within the antigen sequence, resulting in higher molecular weight bands on SDS-PAGE gel. Treatment with PNGase removed these modifications. SCT 1 and 27, lacking NxT motifs in their antigen sequence, showed no size shift. **c.** Discovery of novel CD4 TCRs from healthy donor PBMCs using the CEFT SCT library. Representative flow cytometry plots show tetramer-positive cells captured with SCTs A-E from the 23-element library.

**Extended Data Figure 2.**
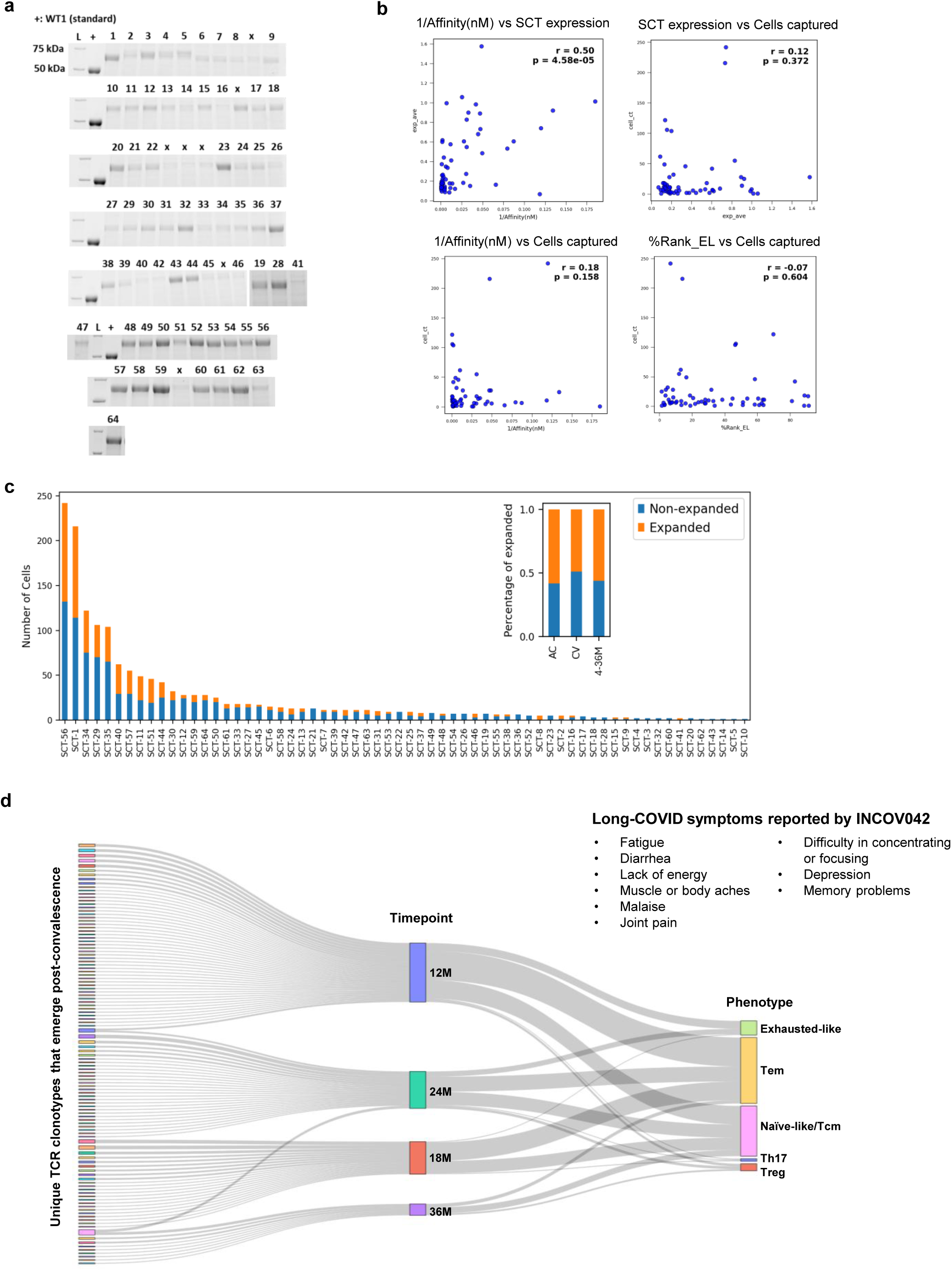
Systematic screening across SARS-CoV-2 RBD identifies large-scale antigen-specific CD4+ T cells. **a.** SDS-PAGE analysis of SCT expression for the SARS-CoV-2 RBD library. The SCT library covers the entire RBD region (1-46) and selective regions from non-RBD spike protein (47-59), nucleocapsid protein (60-62), and membrane protein (63-64). +, protein standard for quantification of expression level; L, molecular weight ladder; x, insufficient yield for downstream analysis. **b.** Correlation between SCT expression levels, quantity of CD4+ T cells captured, and class II peptide-MHC binding predictions. **c.** Clonal expansion patterns of CD4+ T cell responses across SARS-CoV-2 antigens and timepoints. AC-acute infection, CV-convalescent phase. **d.** Alluvial plot depicting newly emerged SARS-CoV-2 specific CD4+ T cell clonotypes in donor INCOV042 post-recovery. Clonotypes from five phenotypes are tracked across 12M-36M timepoints.

**Extended Data Figure 3.**
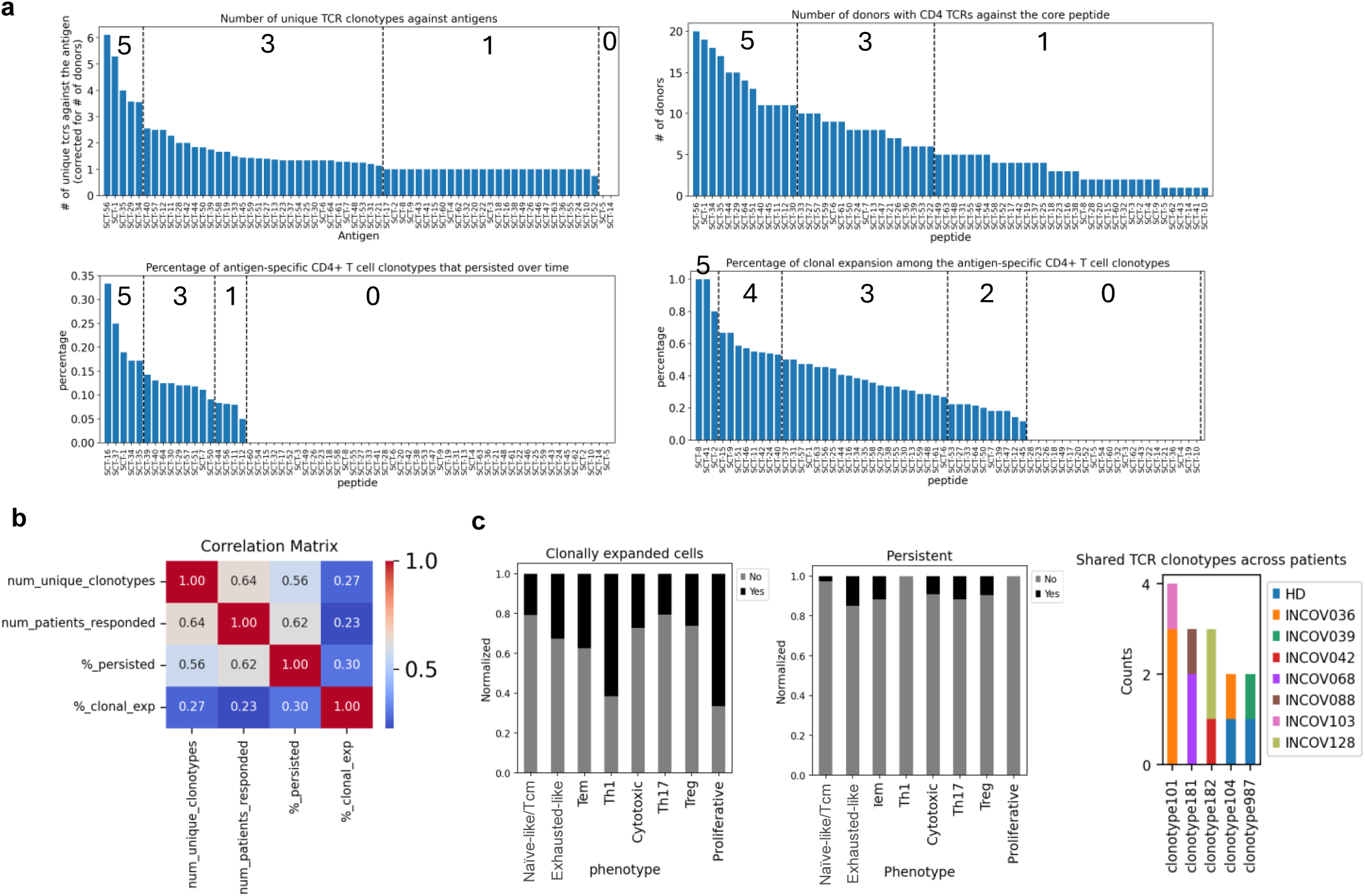
Multi-modal profiling enabled by the SARS-CoV-2 SCT library supports in-depth analysis of fundamental immunological properties. **a.** Ranking and scoring metrics used for evaluating immunogenicity. Antigens were grouped by log_2_-transformed values and assigned scores from 0 to 5 for each property. **b.** Spearman correlation matrix of four immunogenicity properties. All correlation coefficients are statistically significant (p < 0.05). **c.** Clonal expansion and persistence of SARS-CoV-2-specific CD4+ T cells across phenotypes and public TCR clonotypes discovered across patients.

**Extended Data Figure 4.**
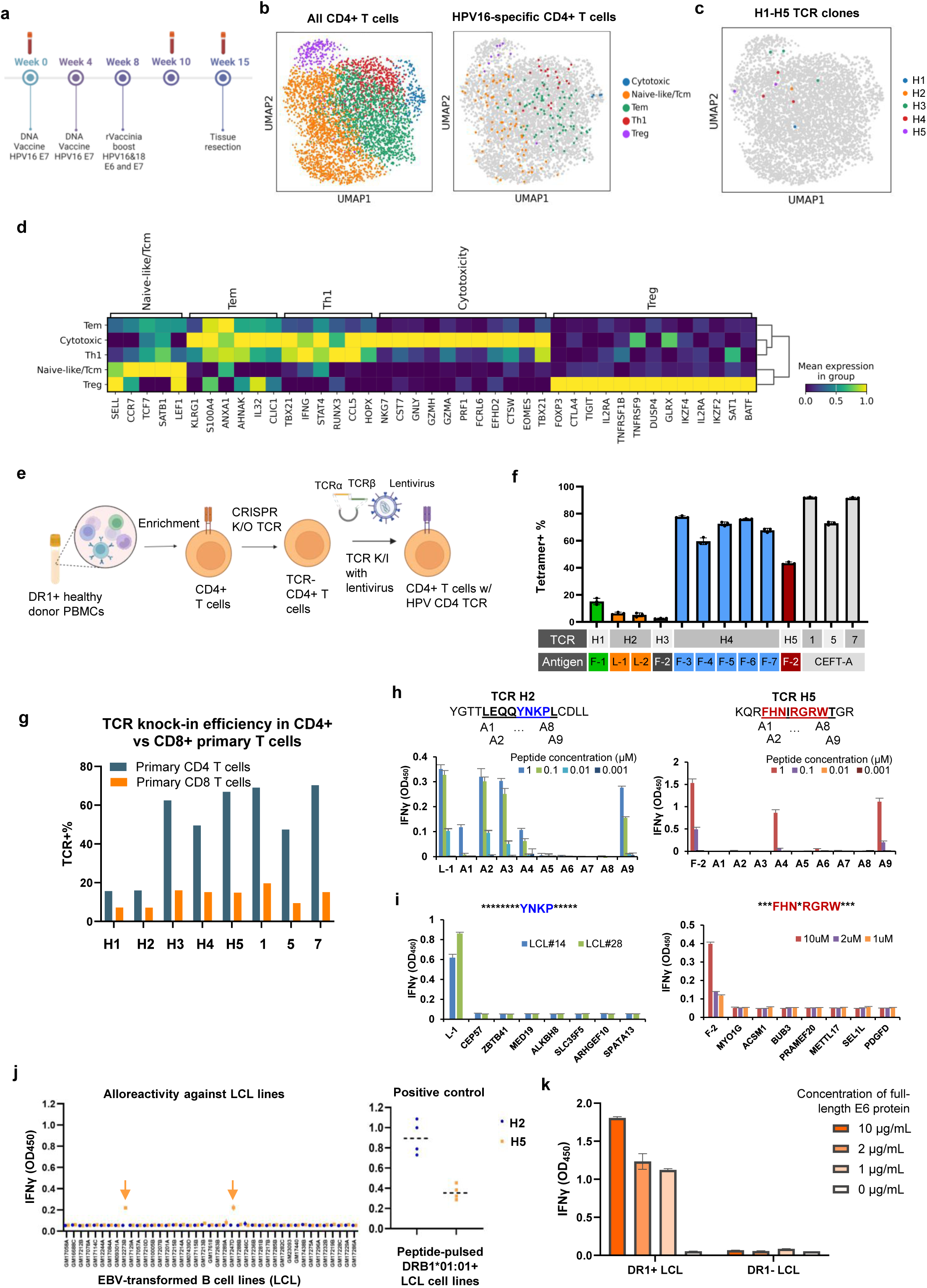
Comprehensive assessment of functional heterogeneity of HPV16-specific CD4 TCRs identified through whole-proteome screening. **a.** HPV-16 therapeutic vaccine clinical trial design. Patients received vaccines targeting E6 and E7 proteins at Week 0, 4, and 8. PBMCs were collected pre-vaccination, at Week 10, and at Week 15 post-series completion. **b.** *left,* UMAP clustering of CD4+ T cells from PBMCs of HPV16+ patients with persistent precancerous lesions. *right,* HPV16-specific CD4+ T cells identified by SCT-dextramers. **c.** Highlighting of H1-H5 TCR clones among HPV16-specific CD4+ T cells. **d.** Expression of signature gene markers for defining phenotype clusters. Mean expression levels were normalized across groups. **e.** Engineering of primary CD4+ T cells with HPV-specific TCRs. CD4+ T cells were enriched from haplotype-matched PBMCs, endogenous TCRs were knocked out using CRISPR, and HPV TCRs were transduced via lentivirus. **f.** Validation of TCR specificity by tetramer binding *(n=3)*. **g.** Transduction efficiency of HPV-specific CD4 TCRs into primary CD4+ versus CD8+ T cells. **h.** Alanine scanning to identify key TCR recognition motifs. H2- and H5-TCR-transduced CD4+ T cells were cocultured with DR1+ K562 cells pulsed with alanine-substituted 9-mer core peptides (A1-A9) across 0.001 – 1 μM. IFNγ secretion was measured (OD_450_). **i.** Cross-reactivity analysis against human self-antigens. H2- and H5-TCR-transduced T cells were co-cultured with LCLs pulsed with peptides identified from BLAST search and motif scans. H2 TCR used a single 1 µM dose; H5 TCR was dose titrated at 1, 2, and 10 µM. **j.** Alloreactivity screen. H2- and H5-TCR-transduced CD4+ T cells were tested against 41 LCL lines representing 92 distinct class II HLA alleles. DRB1*01:01-positive LCLs (*n=4*) pulsed with E6_91-107_ or E6_129-142_ served as positive controls. Alloreactivity of H5 to HLA-DRB1*13:05-positive LCLs is marked with orange arrows. **k.** Confirmation of H2 TCR reactivity to naturally processed E6 antigen. DRB1*01:01+ LCL (GM17281B) was titrated with full-length E6 protein (0-10 μg/mL), co-cultured with H2-TCR-transduced CD4+ T cells, and evaluated for IFNγ secretion. A DRB1*01:01-LCL (GM12244A) served as a negative control.

